# *Staphylococcus haemolyticus* Population Genomics Provides Insights into Pathogenicity and Commensalism

**DOI:** 10.1101/2025.11.08.687178

**Authors:** Heather Felgate, Lisa Lamberte, Leonardo de Oliveira Martins, Antia Acuna-Gonzalez, Janet E. Berrington, Jonathan A. Chapman, Dheeraj Sethi, Christopher J. Stewart, Ad C Fluit, Matthew K. O’Shea, Jorunn Pauline Cavanagh, Lindsay Hall, Willem van Schaik, Mark A Webber

## Abstract

*Staphylococcus haemolyticus* is a common commensal bacterium but also an opportunistic pathogen, frequently implicated in bacteraemia and sepsis in preterm neonates and immunocompromised patients.

Despite its clinical relevance, relatively little is known about the population structure of *S. haemolyticus* and how this relates to its ability to colonise humans or cause disease. In this study, we analysed commensal and clinical strains isolated from neonates and adults from 20 countries between 1957 - 2022. Whole genome sequencing of these isolates, combined with publicly available data, generated a comprehensive dataset of 986 genomes. This enabled us to characterise the species population structure and track the distribution of antimicrobial resistance (AMR) and virulence determinants.

Our analysis revealed a highly diverse genome structure, with multiple phylogenetic groups showing distinct associations with commensalism or pathogenicity. We observed extensive variation in mobile genetic elements, prophages, and AMR genes, alongside increased carriage of plasmids and AMR genes over time. Furthermore, genes lined to metal homeostasis, detoxification, and oxidative stress tolerance were differently abundant between commensal and clinical isolates.

This work provides the most detailed view to date of *S. haemolyticus* diversity, its evolutionary dynamics, and the genetic factors that may underpin the transition from commensal to pathogen.

## Introduction

*Staphylococcus haemolyticus* is a prominent member of the coagulase negative Staphylococci (CoNS). While commonly a human commensal, it is also an important opportunistic pathogen. *S. haemolyticus* infections frequently affect immunocompromised patients in intensive care and preterm neonates in neonatal intensive care units (NICUs) where they can cause bacteraemia and sepsis [1–5]. A recent analysis of CoNS isolation trends in England reported over 4000 annual isolations of *S. haemolyticus* from sterile sites, although this is likely an underestimate as more than 36% of CoNS isolates were not speciated. Notably, antibiotic resistance rates were higher for *S. haemolyticus* than for other CoNS with the highest prevalence of resistance observed for 6 of the 9 antibiotics tested [6]. The study also identified a bimodal age distribution in clinical isolates, with incidence peaks in children under five years (where rates were highest) and adults over 65 years (where absolute case numbers were greatest) [6, 7].

Preterm infants often experience a highly medicalised start to life, particularly those admitted to NICU, where invasive procedures such as line insertion are common and increase the risk of bloodstream infection. An underdeveloped immune system and immature skin barrier, combined with prolonged hospitalisation and antibiotic exposure, can profoundly impact early life microbiome diversity [8]. Globally, Late Onset Sepsis (LOS, defined sepsis occurring ≥72 hours post birth) was implicated in an estimated 946 cases per 100,000 live births in 2021, with a mortality rate of 16.4% [9]. CoNS are among the most frequently isolated organisms in LOS [10–13]. In some settings *S. haemolyticus* strains have become endemic and treatment is complicated by the high prevalence of multidrug resistance (MDR) typically seen in this species [14]. LOS significantly increases hospital stay, invasive interventions, and antibiotic use, all of which negatively affect long-term health outcomes in neonates [10–12].

In adults, *S. haemolyticus* also causes bacteraemia and has been associated with meningitis following traumatic brain injury or neurosurgery, where the central nervous system barrier is compromised. However, infections can occur across a wide patient spectrum [15]. Treating *S. haemolyticus* infections is challenging, particularly as biofilm-forming strains are linked with prosthetic joint infections, ventriculo-peritoneal shunts, and other nosocomial infections [1, 2, 14, 16].

*S. haemolyticus* strains commonly show resistance to β-lactams, (e.g. penicillin, ampicillin) gentamicin, erythromycin and ciprofloxacin [2, 4, 6, 16–20]. Reports indicate that up to 88-100 % of clinical isolates are MDR [3, 5]. The *mecA* gene, with reduced affinity for β-lactams, is frequently carried, although its presence does not always confer phenotypic resistance [17, 18]. While resistance to vancomycin remains rare, intermediate resistance is increasingly observed in CoNS in nosocomial environments [4, 18, 19, 21].

We recently assembled and characterised panels of CoNS from diverse sources [22–25]. In this study, we complied a large collection of *S. haemolyticus* isolates from neonatal and adult blood cultures, as well as from asymptomatically colonised individuals. This allowed us to investigate the genomic diversity of this opportunistic pathogen. In this study we aimed to identify the population structure of the species, identify genetic determinants associated with infection versus carriage, to characterise the AMR reservoir present in *S. haemolyticus* and explore the likely evolutionary history of this important species.

## Results

### Study population and distribution

The panel of isolates was assembled by combining existing collections of *S. haemolyticus* isolated from the UK (Norwich, Newcastle and Birmingham) and Europe (Norway, Germany and the Netherlands) which also included samples from other sites. Isolates were a mixture of clinical specimens and commensal isolates taken from various body sites (Figure 1). This collection comprised of 820 *S. haemolyticus* strains, all of which were sequenced using Illumina technology, with a subset of 36 genomes also being sequenced with Oxford Nanopore Technologies long-read sequencing. We supplemented this dataset with 166 genomes available from NCBI (taxidID 1283) giving a total of 986 genomes.

**Figure 1.**
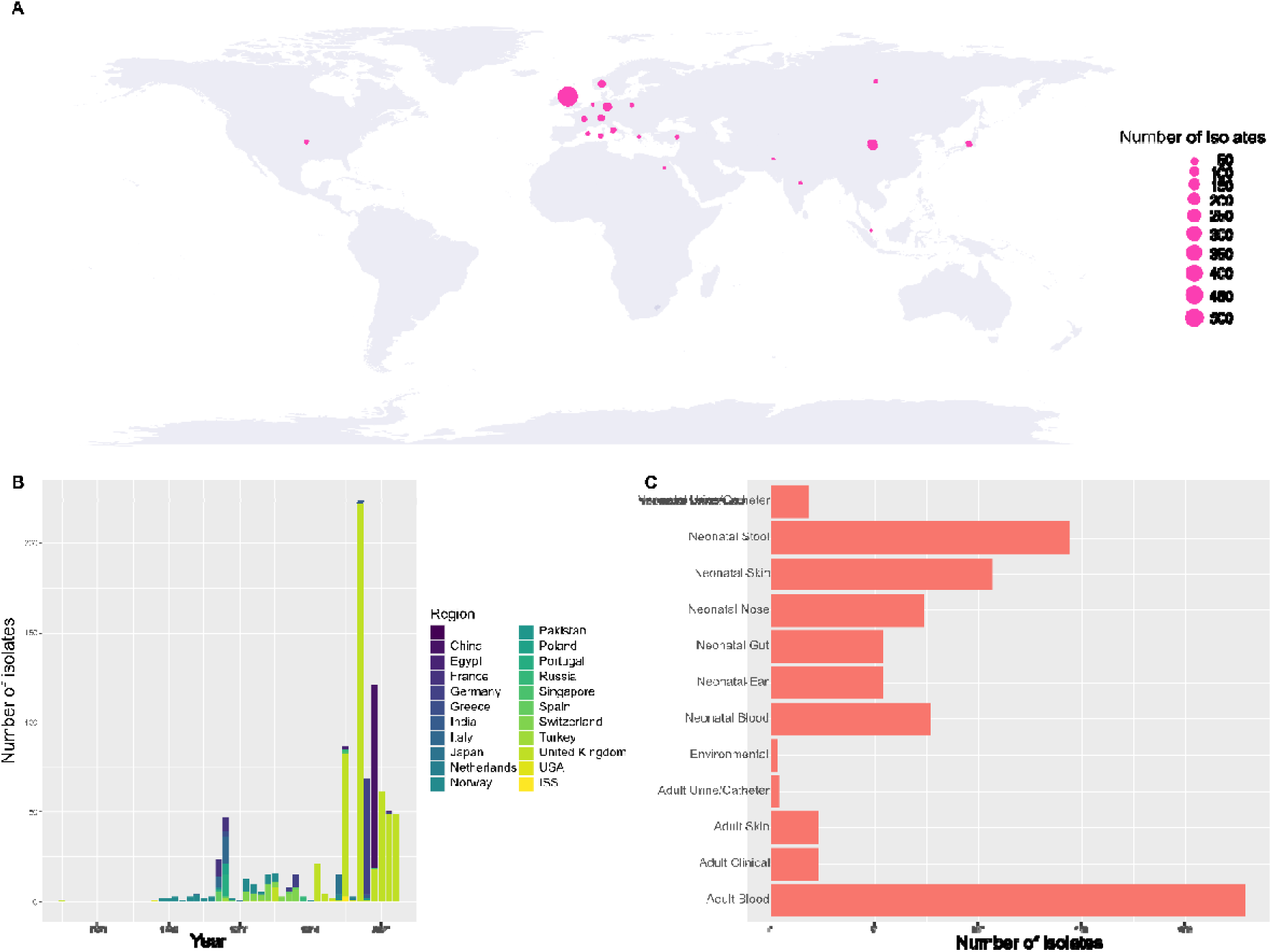
The collection of *S. haemolyticus* collected and its distribution across the globe (A) and year and country of isolation (B). The sources of the collection included adult and neonatal clinical and carriage isolates (C).

The strains in the resulting panel were collected from a total of 20 countries. Most (56.9%) originated from the UK, with 10.5% from China and 8.2% from Germany (Figure 1a and 1b, Supplementary Table 1). Isolates were collected between 1957 and 2022, with the median year of isolation being 2017. Genomes of isolates collected in 1989 or earlier (83) make up 8.4% of all genome sequences in the collection (Supplementary Table 1).

*S. haemolyticus* isolated from neonates accounted for most of the collection (60.2%) with adult clinical samples accounting for 26.1% of genomes (Figure 1c, Supplementary Table 1). Genomes taken from NCBI often had little or no known associated metadata apart from the country of isolation. A small number of isolates (3) were from environmental sampling of the international space station.

### Phylogeny of S. haemolyticus

The average genome size of isolates after assembly was 2.76 Mbp with a GC content of 32.69 %. All genomes were annotated and there was a core genome (present in 99 % of isolates) of 356 genes with a shell genome (present in 75 % of isolates) of 1329 genes.

To understand the genetic diversity of *S. haemolyticus* in the study population we built two phylogenetic trees, one using SNPs from an alignment of the core genome of all isolates using Snippy-core [26] and a core gene presence and absence tree [27] was also built. Both showed a similar structure which revealed a high level of diversity across the species (Supplementary Figure 1 and Figure 2). The number of SNPs between strains in the core genome varied from 3 to 49,937 (with an average of 7116) when compared to a reference genome (chosen as an isolate representing the largest clade with a hybrid genome assembly available). The core gene presence and absence tree divided the collection into five groups, named A-E. The three most common sequence types observed across the population were ST 25 (n=162) which are mostly found in Group D, ST 3 (n= 145) which were found across groups A, B, C and D and finally ST 49 (n=122) which were mainly associated with Group D. This structure was also compatible with analysis of genetic clusters with FastBAPS (Supplementary Table 1) [28].

**Figure 2.**
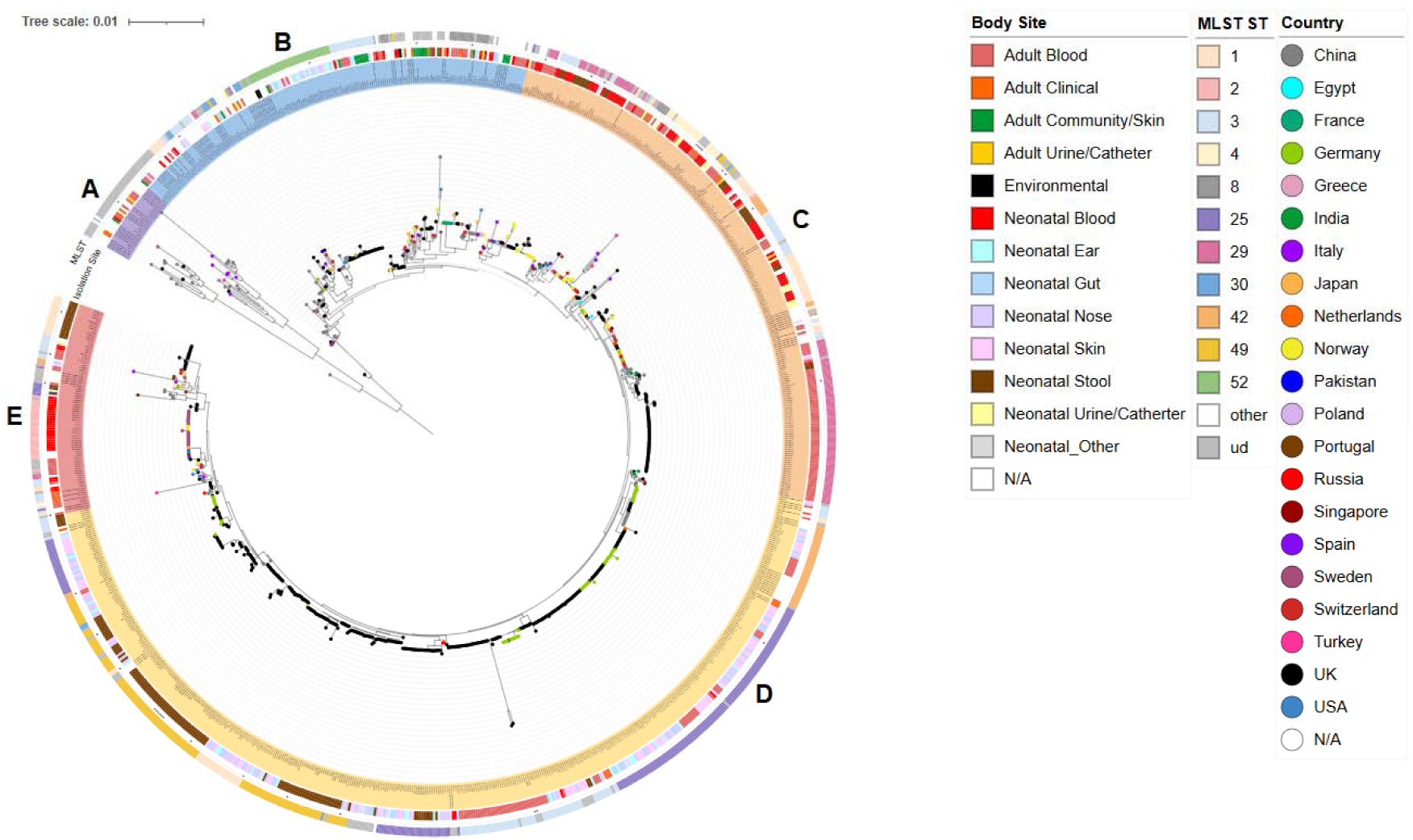
A phylogenetic maximum likelihood tree inferred using IQTree based on a roary core genome alignment [27]. The tree was annotated and visualised in iTOL [29], branch dots represent country, the inner circle represents the 5 predicted major groups (A (red), B (blue), C (orange), D (yellow) and E (red)) the second circle indicates site of isolation (see key). Genomes with additional longread assemblies are indicated by a star next to the third ring and the MLST ST is indicated in the outer most ring.

**Group A (n=32)** This small group is mainly composed of genomes recovered from NCBI (59.4 %) isolated in 2019 with no metadata attached. Isolates with a known source were from adult blood or other clinical specimens (31.25 %) with one neonatal blood isolate also included. This group is the most genetically diverse of *S. haemolyticus* isolates, with an average within-group nucleic acid identity of 96.7% ± 0.08 (Supplementary Table 1) and 90.5% were unable to be assigned an MLST.

**Group B (n=178)** This group contained most of the isolates from adult healthy skin swabs (22/24). It also contained the only isolate from breast milk and the environmental isolates. Blood isolates from adult and neonates accounted for 26 % of this group. Isolates spanned 17 different countries and dated from to 1957 to 2022 (Supplementary Table 1). A total of 24.7 % of isolates had no determined MLST, 25.4 % were assigned ST 52 and 20.2% to ST 3.

**Group C (n= 241)** This group contained a large proportion of neonatal and adult blood isolates (26.5% and 41.4%, respectively, of the group), with very few originating from carriage (13%). Isolates ranged from 1990 – 2022 spanning 16 countries (Supplementary Table 1). The MLST ST mostly observed in the group was ST 29 (41.31 %).

**Group D (n=454)** showed the least genetic diversity between isolates within the group (FastANI 99± 0.01). Isolates were mostly associated with neonatal carriage, accounting for 74.6% of the group. Most of the isolates were from the UK and Germany. Very few isolates associated with neonatal sepsis were in this group (<1%) although 17% were from adult blood. Years of isolation were from 1998-2022 (Supplementary Table 1). In total 153 isolates were assigned ST 25 (36.7%) and 109 isolates were ST 49 (26.14%).

**Group E (n=81)** This group mainly comprised bloodstream isolates. Over 75% of this group were isolates from NICU, with over a third of the isolates found in neonate blood (35.8%), and all 15 isolates from adults were also from blood culture. Isolates from this group spanned 12 countries, 28.4% being from the UK and 25.9% from Sweden, which were involved in a Swedish outbreak (21/37) as described in Westberg, Stegger [14]. Carriage isolates were likely to be from the gut/stool in this group (24.7%). Isolates ranged from 1989-2022 (Supplementary Table 1) and the most associated sequence type was ST 2 (40.6%).

### Carriage of mobile genetic elements, AMR genes and virulence factors

*S. haemolyticus* are often found to harbour aminoglycoside resistance genes, with the genes *aac*(6’)-*Ie*-*aph*(2’’)-*Ia*, *aad*(6), *aph*(3’)-IIIa, *ant*(4’)-Ib, SAT-4 and *lmrS* being commonly reported [30]. In agreement with this, most isolates (844/986) in the panel did carry aminoglycoside resistance genes. Of these, the majority were in the B group (71%), which contained mostly commensal isolates (and some environmental isolates) with relatively few from infection. More than half of the collection (n=454) carried two or more aminoglycoside resistance genes. Amongst these, the SAT-4 gene was present in 366/986 isolates and *aph*(3’)-IIIa was found in 385/986 of the isolates. These alleles have previously been identified on the same element [31], which provides an explanation for linkage of these genes. Five further isolates (found in Groups C, D, and E) carried *aad*(6), as well as APH(3’)-IIIa, and SAT-4, which reflects the structure of the aminoglycoside-streptothricin resistance gene cluster originally found in *Enterococcus faecium* [31]. A total of 96 isolates harboured 4 different aminoglycoside genes, with 83% of these being in the C group, most of which were isolated from adult blood.

The carriage of *mecA* was almost ubiquitous in nosocomial isolates, there were only 43/392 clinical isolates (adult and neonatal nosocomial isolates) that did not carry *mecA,* 54.7% of which were from Group B (spanning 9 different countries). This is likely to reflect the known dynamic nature of the cassettes carrying *mecA* which are capable of spontaneous excision from the chromosome. In total 91 isolates contained no known SCC*mec* cassette, with the majority being in the B group, this coincides with the 86 isolates having no *mecA* gene (48% of the B group).

Analysis of the SCC*mec* cassettes harboured by *S. haemolyticus* identified 12 types of known SCC*mec* cassettes, with the most abundant version being Type V which accounted for more than a third of the total found (Table 1), [32]. The type V system only contains *mecA* and no other resistance genes [32]. The second most prevalent cassette was the type IX cassette, which accounted for nearly 19% of the cassettes present. SCC*mec* type IX is known to contain the *ars, cop* and *cad* genes as part of the cassette [33]. Most of these were found in the C group which is strongly associated with sepsis.

**Table 1.** Frequency and type of SCC*mec* cassettes identified in the collection.

| **SCCmec** | **A** |  | **B** |  | **C** |  | **D** |  | **E** |  |
| --- | --- | --- | --- | --- | --- | --- | --- | --- | --- | --- |
| **Type** | % | n | % | n | % | n | % | n | % | n |
| IB | 22.58 | 7 | 17.98 | 32 | 2.07 | 5 | 0.66 | 3 | 0.00 | 0 |
| IIA | 0.00 | 0 | 0.56 | 1 | 0.41 | 1 | 0.22 | 1 | 0.00 | 0 |
| IIIA | 0.00 | 0 | 2.25 | 4 | 7.88 | 19 | 7.27 | 33 | 14.81 | 12 |
| IVA | 3.23 | 1 | 16.29 | 29 | 24.07 | 58 | 3.52 | 16 | 4.94 | 4 |
| V | 3.23 | 1 | 6.18 | 11 | 6.64 | 16 | 63.44 | 288 | 28.40 | 23 |
| VI | 0.00 | 0 | 4.49 | 8 | 0.00 | 0 | 0.00 | 0 | 0.00 | 0 |
| VII | 3.23 | 1 | 1.12 | 2 | 0.00 | 0 | 0.66 | 3 | 1.23 | 1 |
| VIII | 9.68 | 3 | 3.93 | 7 | 1.66 | 4 | 2.64 | 12 | 1.23 | 1 |
| IX | 9.68 | 3 | 17.42 | 31 | 34.85 | 84 | 9.25 | 42 | 33.33 | 27 |
| X | 3.23 | 1 | 4.49 | 8 | 9.96 | 24 | 3.96 | 18 | 1.23 | 1 |
| XII | 0.00 | 0 | 1.12 | 2 | 0.00 | 0 | 0.00 | 0 | 0.00 | 0 |
| XIII | 6.45 | 2 | 5.62 | 10 | 5.39 | 13 | 3.08 | 14 | 9.88 | 8 |
| None | 41.94 | 13 | 18.54 | 33 | 7.05 | 17 | 5.29 | 24 | 4.94 | 4 |

% refers to the percentage of each group carrying the relevant SCCmec variant, and n refers to the corresponding number of strains.

Other genes associated with resistance that were less frequent were genes for chloramphenicol resistance. Chloramphenicol acetyl transferase (CAT) was mainly caried in isolates in the C group from Norway and Switzerland (n= 18 and 8 respectively). Of the isolates with CAT (n=68), 76.5% were from neonatal or adult blood/catheter samples.

The *tcaA* gene is associated with teicoplanin resistance and some SNPs in *tcaA* have been shown to cause changes in cell wall composition and increase tolerance to glycopeptides [34]. We identified several SNPs in *tcaA* in the panel. A total of 60.9 % (70/115) of neonatal blood isolates and 88.2 % (15/17) of isolates from neonatal catheters contained various SNP(s) in this gene. Under half (41.9 %) of the adult blood isolates had SNPs in *tcaA* (Supplementary Table 3).

There were statistically significantly fewer AMR genes observed in groups A (mainly consisting of NCBI genomes) and B (mainly carriage and environmental isolates) with ∼7 genes per isolate compared to the other 3 groups where strains had an average of over 10 AMR genes (Figure 3, Supplementary Figure 2). There was a maximum of 20 AMR genes in any individual strain.

**Figure 3.**
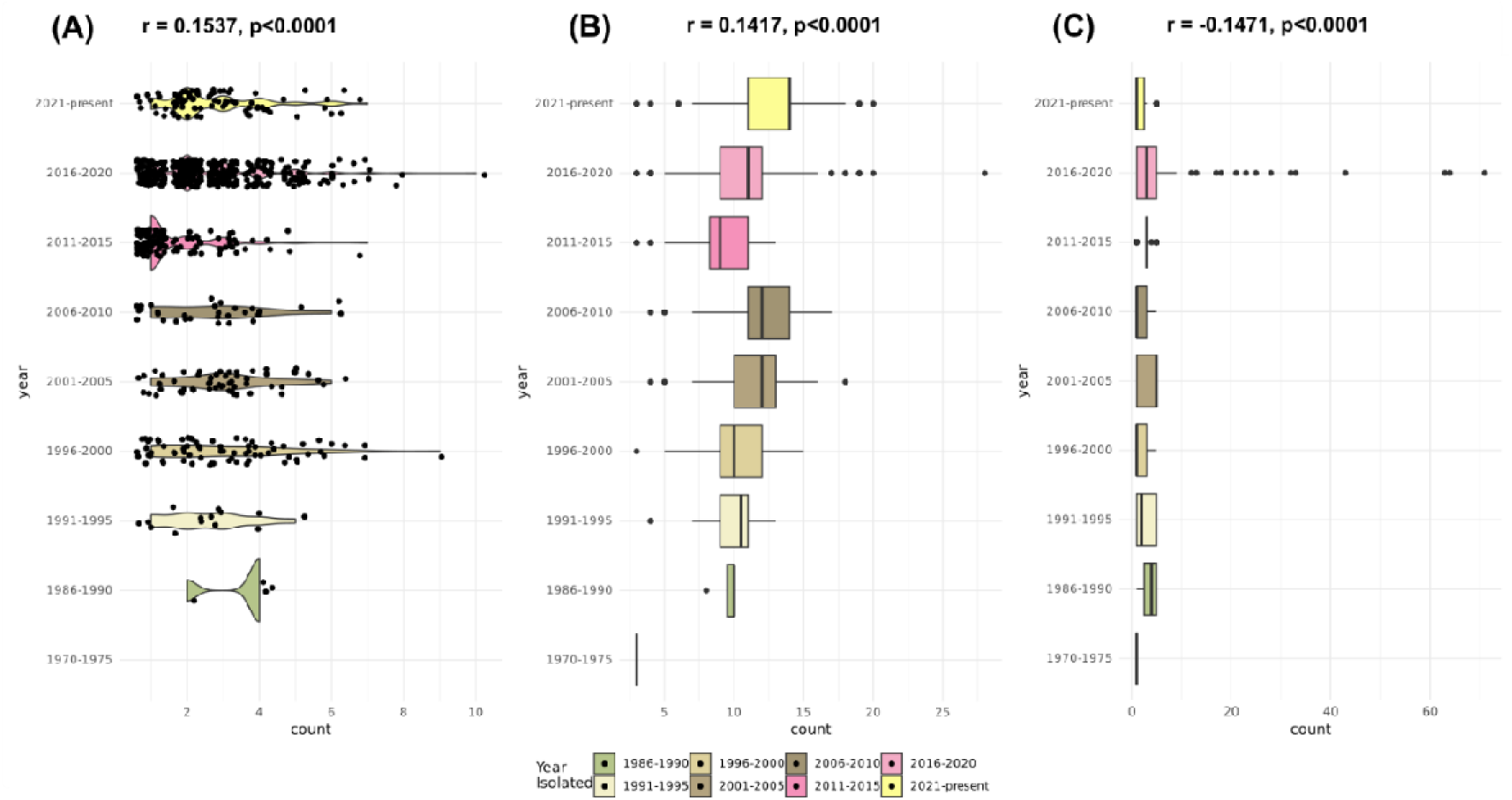
Diversity of plasmid replicons and genes associated with antimicrobial resistance. Violin plots show the number of plasmid replicons detected per *S. haemolyticus* genome per year group of isolation (rho = 0.15, p < 0.0001) (A). Boxplots indicate the number of antibiotic resistance genes (rho = 0.14, p < 0.0001), and (B) virulence factors (rho = –0.15, p < 0.0001) (C) detected across the same groups. Statistical significance was assessed using Spearman’s rank correlation, testing for monotonic trends over time.

The number of AMR genes identified in strains correlated with the date of isolation and increased over time. This was found to be a weak but statistically significant positive correlation between the year of isolation and the count of AMR genes (r = 0.1417, p < 0.0001) (Figure 3). Conversely, there was no increase in the carriage of virulence genes over time and in fact a small but statistically significant negative correlation was seen between the year of isolation and the count of VF genes (r = -0.1471, p < 0.0001).

Virulence Factors (VF) were identified across the collection from the virulence finder database [35]. Groups D and E had the highest number of VF genes per genome with >4 per strain on average compared to ∼2 in groups A, B and C (Figure 3). This was reflected in the maximum number of VFs in a single isolate being 12 being amongst groups A, B and C. Whereas in groups D and E the maximum number of VFs per isolate was 71 and 64, respectively (Supplementary Figure 2). Virulence factors associated with capsule production were more present in groups D and E, compared to the other groups (Supplementary table 2), specifically the *cap8E* and *cap8G* genes. Group D strains also carried more genes involved in adherence (*ebp* and *ica* genes) and iron acquisition (*isd* genes, Supplementary table 2).

Plasmids can significantly influence the structure and evolution of bacterial genomes. To assess plasmid diversity in our *S. haemolyticus* collection, we investigated the presence of plasmid replicon initiation proteins (*rep* genes) as indicators of plasmid presence [36] (Figure 3, Supplementary Data 4). Most strains contained at least one *rep* gene indicating a widespread prevalence of plasmid carriage and we detected a range of 1 to 10 *rep* genes per genome (Figure 3), no replicons were though detected in *S. haemolyticus* NCTC 11042, isolated in 1975 from an adult skin sample in the UK. In total, 55 distinct variations of *rep* genes were observed across the dataset (Supplementary Table 4), highlighting the substantial diversity of plasmid replicons within our *S. haemolyticus* collection. *S. haemolyticus* NW30008, isolated in 2017 from a neonatal nose sample in the UK, carried the highest number of *rep* genes (10).

Plasmid replicon counts varied across time periods, with a general shift toward higher counts in more recent years (Figure 3). Earlier time periods (e.g., 1986–1995) were dominated by isolates carrying fewer replicons (typically 4–5), while later periods (e.g., 1996–2020) showed broader distributions with more isolates carrying 7–10 replicons. The most recent period (2021–present) showed a slight reduction in plasmid count compared to the 2016–2020 period. While distributions overlapped considerably, there was a small but statistically significant positive correlation between year of isolation and replicon count (rho = 0.1537, p < 0.0001), suggesting a modest upward trend. We also explored whether plasmid replicon counts were associated with the five phylogenetic groups identified in our dataset. A Spearman’s rank correlation test yielded a non-significant result (rho = –0.0308, p = 0.3928), indicating no meaningful association between replicon count and phylogenetic grouping.

Haemolysin III (*hla*) (annotated as *yqfA* in Staphylococci) [*37*], was almost ubiquitous, being found in 960/986 isolates. Another haemolysin originally identified in *S. epidermidis, yidD*, was also highly prevalent and found in 99.4% of isolates [38]. Most isolates (97%) contained both genes, with only 3 isolates having no known haemolysin genes.

### Presence and diversity of phage in *S. haemolyticus* genomes

Putative phage DNA was identified in 217 out of 986 (22%) of the genomes examined, with the number of separate phages detected per genome ranging from 1 to 30, with a median of 2. The proportion of the genome occupied by phage DNA also varied widely, ranging from 0.06% to 21.5%, with a median of 8%. Strain CUHP01, isolated in 2008 from a neonatal blood sample in Germany, harboured the highest number of phages (30) (Supplementary Tables 1 and 5).

Phage DNA within the collection exhibited considerable diversity, with phage genome island sizes ranging from 1.4 kb to 552 kb. The largest predicted phage genome size was observed in strain CUDE01, isolated in 2001 from a neonatal blood sample in Norway (Supplementary Tables 1 and 5).

Regarding temporal changes in phage composition, no clear trend was observed across the five-year intervals, as substantial variation was present within each group (Figure 4). These findings suggest a persistent and diverse presence of phages has been present within the *S. haemolyticus* population over time.

**Figure 4.**
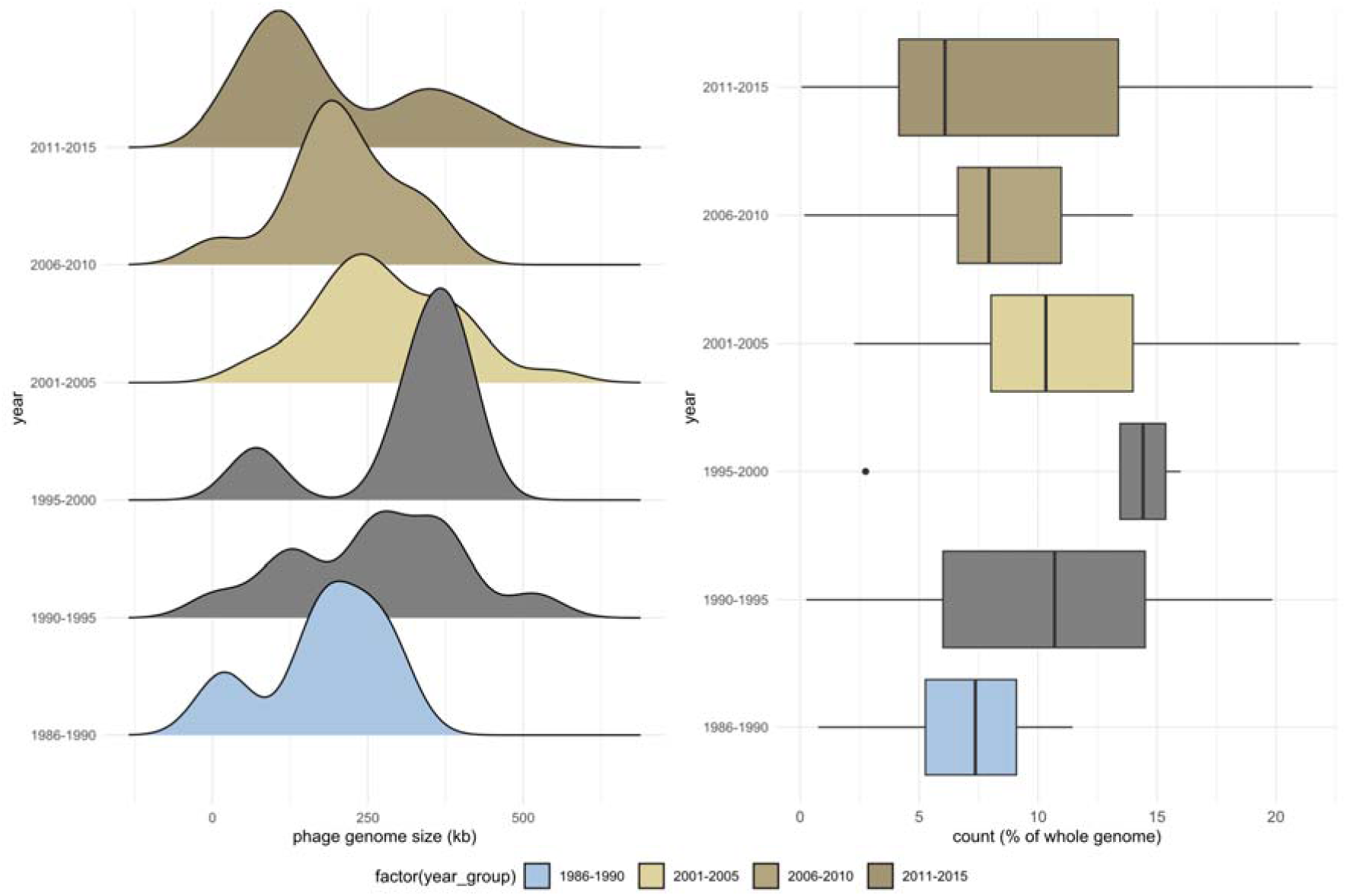
Phage diversity. Density plots (left half) show the distribution of concatenated phage sequence sizes (shown as average per genome) per year group of isolation of *S. haemolyticus* isolates. Box plots (right) show the percentage of genome sequences composed of predicted phage DNA per year group of isolation.

### Associations of genes with phenotypes of interest

To find genes associated with specific traits, e.g. isolation from blood rather than another body site, we used scoary to identify pairwise comparisons (potentially significant associations were considered where adjusted p-values were ≤ 0.0001) [39]. Genes with a significantly different abundance between gut carriage (stool isolates) and the rest of the collection included those involved in metal sequestering and tolerance. A series of genes were less abundant in stool isolates including *arcA, arsC, czrA, fer, fur, feoA* and *mntH*. AMR genes associated with stool carriage were *fusB, mecR, mgrA* and *tetK* (Supplementary Table 3).

Two genes, *hemY* and *arsB* were over-represented in clinical isolates compared with the rest of the collection, *arsBC* is associated with the SCC*mec* Type IX cassette so over-representation in isolates from a clinical source agrees with the high prevalence of this cassette in sepsis isolates described earlier [40]. HemY is a protoporhyrinogen oxidase involved in heme biosynthesis which is known to be important *in vivo*.

Comparison of isolates from neonatal and adult blood cultures found four genes strongly associated with isolation from neonatal blood, these were *pckA* (phosphoenolpyruvate carboxykinase), *menC* and *menE* (involved in o-succinylbenzoate synthesis), and *hemY*.

### Predicted evolutionary history of *S. haemolyticus*

*S. haemolyticus* was first described in 1975, by Kloos and Schleifer [41]. The ancestral evolution of *S. haemolyticus* is uncertain but analysis of mutations in the genome can infer an evolutionary history of the species [42]. A molecular clock phylogeny (Figure 5) was inferred with TreeTime using the isolation dates and population structure as a guide. This resulted in a structure with a similar pattern to the core genome and SNP alignments, with the groups identified in Figure 2, clustering in a similar pattern although groups A and B are not clearly differentiated in this analysis. The predicted tree suggests several major evolutionary steps where groups of *S. haemolyticus* have diverged with the first predicted branching occurring around 1550 where isolates in groups A and B split before subsequent divergence of groups C, D and E (Figure 5). The majority of *mecA* negative isolates are in the B group suggesting that methicillin resistance was acquired independently by the other groups, and that this occurred after the first major branching event detected.

**Figure 5.**
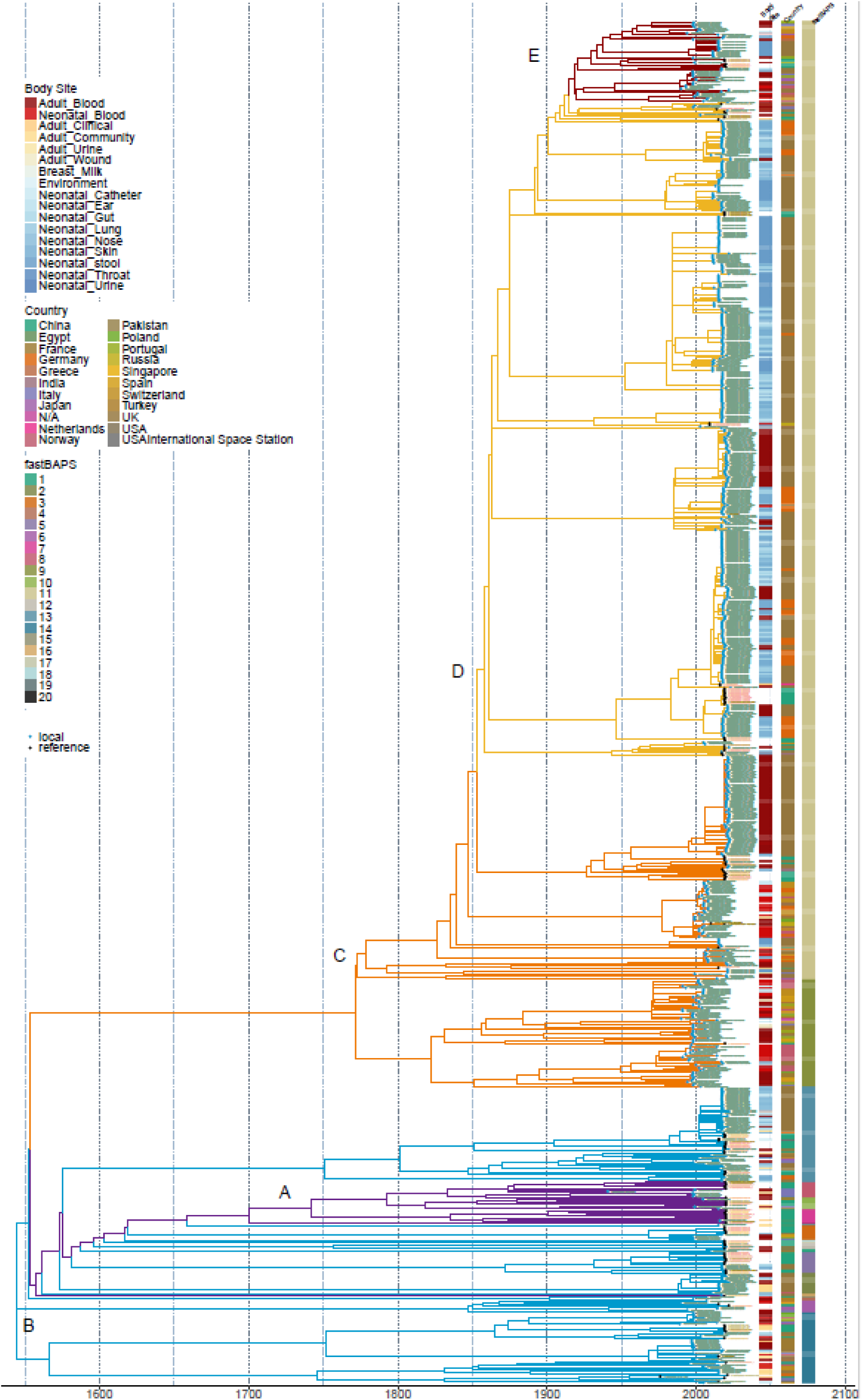
Inferred phylogeny based on molecular-clock analysis. TreeTime [v0.11.2] [42] was used to infer ancestral relationships between strains and predict a phylogeny. Colours of branches refer to the groups previously inferred from the ML tree based on the core genome alignment (Figure 2). Country, source and FastBAPs group numbers are shown as colour blocks (See Legend and supplementary Table 1).

Group D which largely contains commensal isolates is predicted to have evolved from group C where a higher proportion of isolates were pathogenic. This branching is predicted to have occurred in approximately 1857. Later, group E is predicted to have branched from group D (around 1918, Figure 5). Isolates from group E were mainly from blood. This description suggests changes in pathogenic potential have occurred with two successive groups (C and E) having a higher potential to cause bloodstream infection with the intermediate group D containing isolates more likely to be associated with carriage than disease.

## Discussion

Our analysis shows significant diversity between *S. haemolyticus* isolates with a core genome of just over 350 genes and a much larger accessory genome. Based on analysis of the core genome the population is split into multiple distinct groups. Commensal isolates were present in all the major groups, and skin isolates were closely related to those obtained from stool. The predicted population structure is similar to that seen in other studies of nosocomial isolates although our number of genomes is higher than in prior work [39]. Previous research on NICU associated *S. haemolyticus* have reported that isolates involved with neonatal sepsis are largely clonal although most prior studies have focussed on characterising isolates from individual settings. Our analysis suggests that diverse strains of *S. haemolyticus* can be pathogenic. Isolates from previous research on the clonal expansion of *S. haemolyticus* in NICUs by Cavanagh, Hjerde [4] and Westberg, Stegger [14] were present in our collection and found in three distinct groups, E, B and C (all of which were associated with sepsis). Different groups of strains were associated with neonatal and adult sepsis suggesting that whilst pathogenic potential is shared by isolates across the population structure, some groups (e.g. group D) were more commonly associated with commensalism than disease. This highlights that pathogenicity is polyphyletic and likely driven by accessory genome content and environmental context rather than a single hyper-virulent lineage.

None of our *S. haemolyticus* isolates were from adult stool samples although a large proportion were from the gut of neonates in NICU. Various staphylococci are commonly isolated from the gut in neonates [22, 23, 43] and for some species it has been suggested that gut carriage can be a reservoir for infection at other body sites. For example, gut carriage of *S. capitis* strain NRCS-A is linked with sepsis as isolates from stool and bloodstream infections have proved to be indistinguishable [23]. Similarly, *S. aureus* and *S. epidermidis* are common colonisers of the infant gut, particularly from babies in NICUs [37, 43]. Whether carriage of *S. haemolyticus* in the gastrointestinal tract of neonates directly seeds infection at other sites is uncertain although we recently showed a very high abundance of *S. haemolyticus* in the guts of preterm babies in the first 10 days of life and identified a dominant clone of ST49 in samples from neonates in UK NICUs [44]. In this study with a larger panel of isolates we also observed that ST49 was commonly isolated from stool or as a commensal from other body sites from neonates within UK NICUs (122 ST49 strains were almost all from UK NICUs). These isolates were largely in group D which is consistent with this clone being adapted to commensalism as few isolates in this group were associated with disease. Interestingly, group D also carried capsule and adherence genes (e.g. *cap8, ebp, ica*), which may favour colonisation and persistence rather than invasion, reinforcing the idea that virulence is context dependent.

### Carriage of AMR genes

The most genetically distant groups (A and B) contained fewer virulence factors and AMR genes per genome than the other groups, such as having no aminoglycoside resistance genes compared to the remainder of the population. The proposed evolutionary tree (Figure 5) suggests these groups branched off from a common ancestor and have evolved separately to the other groups which contain isolates more associated with nosocomial carriage and infection. Groups A and B also had relatively low numbers of isolates with an SCC*mec* cassette, suggesting that the other lineages acquired SCC*mec* cassettes separately and after the initial predicted divergence (Figure 5).

Resistance to methicillin (and other β-lactams) and aminoglycosides is a key feature known to be common in isolates of *S. haemolyticus*. Various SCC*mec* cassettes have been defined, [45] and we identified a range of SCC*mec* types in our panel although the types and prevalence differed between the groups. Over 90% of all isolates contained a SCC*mec* cassette with the most common being type V, which is consistent with previous reports [32, 45, 46]. Interestingly, and unlike some other species including *S. capitis* we were unable to identify any CRISPR-Cas systems within the SCC*mec* cassettes [23] in any isolates. The A and B groups which are predicted to differ in ancestry from the other groups had the lowest SCC*mec* carriage (19 % and 42 % respectively) suggesting separate introductions of cassettes into different branches of the phylogeny. Previous work by Rolo, Worning [47] postulates that a *S. aureus* clone containing the SCC*mec* III cassette emerged in the 1960 s in Europe. Our analysis suggests multiple, independent acquisitions of the cassette within *S. haemolyticus*. Defining a timescale for these events is difficult given the mobile nature of the element and areas of high recombination within the cassette [48].

Aminoglycoside resistance is known to be common in *S. haemolyticus* isolates and a diverse set of aminoglycoside modifying enzymes (AMEs) were seen in the panel with over 85% of isolates carrying at least one AME, and over half the collection having 2 or more AMEs. These were not over-represented in clinical isolates and were most common in group B, which had a high proportion of commensal isolates. However, a smaller number of isolates carried 4 AMEs which were mainly clinical isolates from group C. This data shows that AMEs are very common in *S. haemolyticus* and explains the high prevalence of aminoglycoside resistance in this species, it is possible that high-level resistance mediated by carriage of multiple AMEs provides a benefit in infection.

Chloramphenicol resistance was rare and geographically restricted, but its presence in bloodstream isolates suggests plasmid-mediated traits can still influence pathogenic potential. Chloramphenicol resistance was only seen in a minority of isolates in this study, from Scandinavia. The CAT gene is carried on small plasmids (∼3.9 kb) proposed to have originated from *S. intermedius* although CAT genes have also been identified in *S. aureus* [49]. CAT genes were found in 68 isolates (supplementary table 2), of which the majority were from neonatal and adult blood samples, suggesting harbouring the plasmid containing the CAT gene increases the pathogenicity of *S. haemolyticus.* Only 4 isolates with CAT were commensal isolates (1 neonatal ear, 3 adult gut), and only one was from the UK.

### Presence of mobile elements

A varied component of phage DNA was identified within the isolate genomes, this was seen in all the groups, and the prevalence of phage DNA did not significantly increase over time suggesting a steady state of phage acquisition and transfer within the population. There was also a large component of plasmids within the strains with most strains carrying plasmid *rep* genes, the number of which per strain varied from 0-10. The prevalence of plasmid carriage increased significantly with time (albeit with a small effect size) suggesting positive selection for plasmids in more recent isolates, potentially associated with antibiotic use and carriage of resistance genes. There was though no significant difference in the plasmid repertoire between the groups of the phylogeny suggesting no barriers to movement between groups are present within the species. Together, these findings indicate that horizontal gene transfer via plasmids and phages remains a major evolutionary force in *S. haemolyticus*, facilitating AMR dissemination and shaping accessory genome.

### Carriage of virulence factors

Haemolysin III was present in most isolates and is essentially a ubiquitous virulence factor in *S. haemolyticus*. *cap* genes are associated with the production of the capsule surrounding the cell and are linked to virulence [50], however, in our analysis we identified the D group (associated with carriage) as harbouring *cap8* (and variations of) as well as having more genes associated with adherence such as *ebpABC*. It has been suggested that *cap8* aids colonisation of new hosts rather than causing pathogenesis [50], *cap5* has been implicated as a virulence factor which was not found in any of our genomes. The observed carriage of *cap8* and absence of *cap5* is consistent with the D group adapting to carriage rather than pathogenicity. Even though the pathogenicity potential of members of this group seems to be lower than others, isolates would still represent a large reservoir of various AMR genes which have the potential to be transferred to other species of staphylococci.

### Genes associated with commensalism versus disease

Clinical isolates were more likely than commensal (stool) isolates to carry a series of genes involved in metal homeostasis, such as arsenic and iron. This suggests that the ability to survive exposure to toxic levels of metals is important in virulence but not required for survival in the gut. Genes involved in haem biosynthesis are often implicated with virulence, and it was found that the neonatal blood isolates often carried *hemY*, suggesting different strains are adapted to neonatal versus adult infection. There were also several AMR genes associated with neonatal stool carriage, this may reflect the high levels of antibiotic use in this cohort (Supplementary Table 3). These associations, while statistically robust, require functional validation to confirm their role in pathogenesis.

### Evolutionary context and implications

The molecular clock analysis suggests early divergence of groups A and B, followed by the emergence of C-E, with group E (largely bloodstream isolates) branching from group D (predominantly commensal). This pattern implies stepwise acquisition of SCCmec and other accessory traits during adaptation to hospital environments. Although precise dates should be interpreted cautiously, these findings align with historical antibiotic use and NICU practices as potential selective pressures.

## Conclusions

We show here that *S. haemolyticus* is a genetically diverse species, with different strains demonstrating distinct properties that have important implications for human health. Carriage *of S. haemolyticus* is ubiquitous across NICU, skin, and gut environments, providing a substantial reservoir of AMR and virulence genes. Our data indicates that isolates have evolved into multiple groups with varying potential to cause disease in neonates and adults, suggesting that not all isolates present an equal risk to health. This work provides a foundation for understanding the evolutionary dynamics of *S. haemolyticus*, mechanisms of bacteraemia, and the dissemination of AMR within this species. However, our study is limited by sampling bias (with overrepresentation of neonatal isolates and UK/European sources), incomplete metadata for public genomes, and the absence of phenotypic validation. Future work could address these limitations by incorporating isolates from a broader range of sources and geographies, applying phylogeny-aware genome-wide association studies to refine genotype-phenotype links, and reconstructing the plasmidome to map AMR gene flux. Finally, prospective surveillance using lineage-aware diagnostics could help identify high-risk clones and inform infection control strategies in both neonatal and adult care settings.

## Methods

*Isolation of S. haemolyticus being carried in NICU -* Infants admitted to the NICUs of the Norfolk and Norwich University Hospital (Norwich, UK) or University Children’s Hospital (Lubeck, German) over ten-weeks in 2017 or 2018 were enrolled in this study as recently described [22]. Swabs are taken routinely from babies upon admission and throughout their stays in both sites for MRSA surveillance, duplicate swabs were taken for this study and staphylococci were isolated. In addition, isolates from positive blood, cerebrospinal fluid, urine, and wound cultures were included, if taken during the study period.

The UK unit enrolment was between November 2017 and January 2018, and the German enrolment was from January to March 2018. Amies Charcoal Swabs (Thermo Fisher Scientific) were used to isolate bacteria from all infants on admission and weekly until discharge. Swabs from the ear, nose, axilla, groin and gut were taken and streaked on 5% horse blood agar (Thermo Fisher), prior to incubation at 37 °C for 24 hours and final identification of coagulase-negative staphylococci after sub-culture on mannitol-salt agar (Oxoid, Thermo Fisher Scientific), coagulase testing (MERCK; 75832), and/or MALDI-TOF mass spectrometry (Bruker) as previously described [22]. Isolates were then stored on preservation beads at -80°C (Protect, Technical Service Consultants Ltd) and in 96 deep-well plates in 20% glycerol at -40 °C. The isolates attained from this study were also analysed for their antimicrobial resistance profiles.

### Collection of S. haemolyticus genomes

All whole genome sequences (WGS) used in this study were confirmed to be *S. haemolyticus* using either Kracken2 [v2.1.1] and Bracken [v2.2] for long read sequences or centrifuge [v0.15] [51, 52]. For short read WGS with a centrifuge relative abundance ≥0.1 of non-*S. haemolyticus* were removed from the final collection, likewise any sequences with a FastANI [v1.1] <95% were also removed [53].

*S. haemolyticus* isolates obtained by The University of Birmingham (ARM and BAMBI) from routine blood cultures taken from neonates with possible sepsis, clinical data was anonymised prior to analysis, in this study a single from the NNUH NICU was associated with neonatal sepsis, confirmed as *S. capitis* by the NNUH MALDI-tof (Bruker), but identified as *S. haemolyticus* using centrifuge [v0.15] (short read WGS). Commensal and carriage *S. haemolyticus* obtained from the Quadram Institute were taken from surveillance MRSA swabs from the Norfolk and Norwich University Hospital[22]. Finally, isolates were donated from Newcastle University that were isolated from preterm infant stool samples. Preterm infants (born at <32 weeks gestation) were born or transferred to a single tertiary level Neonatal Intensive Care Unit in Newcastle upon Tyne, UK, and participated in the Supporting Enhanced Research in Vulnerable Infants (SERVIS) study (REC10/H0908/39) after written informed parental consent.

Previously sequenced genomes of *S. haemolyticus* were obtained from collaborators, 56 genomes from Utrecht University from the European SENTRY surveillance program [54], 122 genomes from Cavanagh, Hjerde [4] and 37 genomes from Westberg, Stegger [14] were added to the collection and confirmed to be *S. haemolyticus,* using centrifuge [v0.15], the assemblies were downloaded from the European Nucleotide Archive (PRJEB2705 and PRJEB56240 respectively). Reference genomes and scaffolds available from the NCBI were downloaded using Taxid ID 1283 (n= 129) from the NCBI website (RefSeq and Genbank) and added to the genome collection[55].

*S. haemolyticus* isolates from the BAMBI study that were also added to the collection are available from Genbank (Accession number: PRJNA1105567). Isolates collected and sequenced at the Quadram Institute are available from the SRA, Accession number: PRJNA1160826.

### DNA extraction

From the NICU NAS collection, isolates were grown in 1 ml Brain Heart Infusion (Sigma) broth overnight at 37 °C. Cultures were pelleted and resuspended in 100 µl 0.5 mg/ml lysostaphin (from *Staphylococcus staphylolyticus,* Merck) and incubated at 37 °C for a minimum of 1 hour. DNA was extracted from the lysate with the Quick-DNA Fungal/Bacterial 96 kit (Zymo Research), kit used in accordance with the manufactures guidelines (centrifuge speed used 2040G).

DNA was quantified using the Quant-iT^TM^ dsDNA HS assay (ThermoFisher), the fluorescence was measured on a FLUOstar Optima plate reader set at 480/530 nm (excitation/emission)

### Whole genome sequencing

DNA extracted from isolates were sequenced in accordance to with our previous publication [23]. The pool was run at a final concentration of 1.5 pM on an Illumina Nextseq500 instrument using a Mid Output Flowcell (NSQ® 500 Mid Output KT v2(300 CYS) Illumina Catalogue FC-404-2003) following the Illumina recommended denaturation and loading recommendations which included a 1% PhiX spike in (PhiX Control v3 Illumina Catalogue FC-110-3001). Data was uploaded to Basespace (www.basespace.illumina.com) where the raw data was converted to 8 FASTQ files for each sample.

For long read sequencing up to 400 ng of DNA was incubated with Ultra II End-prep reaction buffer and Ultra II End-prep enzyme mix for end repair. The samples then had native barcodes (NB01-24) the pooled library was loaded on to a primed MinION flow cell [using R9 4.1 chemistry] (Oxford Nanopore Technologies). Base calling was carried out using Guppy [v6.06] (Oxford Nanopore Technologies).

### Genomic and Phylogenetic analysis

Once generated, sequence data was processed via a series of pipelines on a Galaxy instance hosted at the Quadram Institute Bioscience [56]. For short read Illumina FASTQ files were used to assigned a microbial classification and check for contamination for each sample using Centrifuge [v0.15] [52]. Long read MinION FASTQ.QZ sequences were assigned taxonomy using Kraken2 [v2.1.1] and Bracken [v2.2] [51, 57].

Sequences identified as *S. haemolyticus* strains were then used for further analysis, short read Illumina sequences were assembled using SPAdes [v3.12.0] [58], with default parameters applied, and longread MinION sequences were assembled using Flye [v2.9] [59, 60]. Resulting scaffold assemblies were further analysed for quality with QUAST [v5.0.2] [61]. Isolates with less than X10 genomic coverage were omitted from the final collection. Sequences were further assessed for quality using FastANI [v1.1] and isolates with average nucleic identify of >95% were removed from the collection [53]. MLST (Multi-Locus Sequence Typing) ST (Sequence Type) were determined running contig files against the PubMLST sheamolyticus scheme using the MLST tool [62, 63].

The phylogeny of the *S. haemolyticus* isolates was determined using gene presence/absence after creating a tree based on the core gene alignment. Bakta[v] [64] was used to annotate the assembled contigs along with all available *S. haemolyticus* reference genomes [taxidID 1284, n= 129, https://www.ncbi.nlm.nih.gov] (Supplementary Table 1). The GFF3 files were then submitted to Roary [v3.13.0] to determine the core and accessory genomes (using 75% and 90% sequence identity cutoffs from Blastp, 80000 clusters) [27]. A phylogenetic tree was inferred using IQTREE [v 1.6.12] with >1000 bootstrap replicates [65] from the core gene alignment output from roary. TreeTime [v0.11.2] [42] was used to infer molecular clock phylogeny of the collection based on the core genome alignment and date of isolation (Supplementary Tabel 1). A core-SNP tree was also inferred using IQTREE on the core gene alignment created by Snippy-core [v4.4.3] [26] using default parameters and then processed with Gubbins [v2.4.1] [66] to account for any recombination. Snp-Dist [v0.6.3] [67] was used to compare the number of SNPs between the isolates from the core alignment. All trees were visualised and annotated in iTOL [29]. Tree structure was further confirmed using hierarchical clustering tool fastBAPS [v 1.0.8, for R version 4.4.1] [28].

Scoary [v1.6.16] was used to calculate associations from the roary gene presence and absence output to calculate associated genotype and phenotypes [39]. The roary core genome maximum likelihood tree was used in the scoary analysis, genes identified as significantly associated had a best pairwise p-value <0.0001 and with a specificity score of >50. When identifying genes significantly associated with group (Figure 2), no tree was required and the Benjamini-Hochberg adjusted p-value was used <0.001, with a specificity score >50 and sensitivity score of >70. Raw data from scoary can be found in Supplementary Table 6.

The presence of antimicrobial resistance and virulence genes were identified (75% minimum identity, and 85% minimum coverage filters applied) using ABRicate (0.9.7) [68] and the CARD [v3.1.1] and Virulence Finder DataBase [v6.0] [35, 69]. To identify the *SCCmec* cassettes, reference genomes (downloaded from *SCCmec* finder [33])were used to create a BLAST+ blastn [v2.10.1+galaxy0] database [70, 71]. This database was run against the *S. haemolyticus* collection nucleotide assemblies, using megablast [2.10.0] with default parameters.

The association between the number of plasmid replicons, antimicrobial resistance (AMR) and virulence factor (VF) genes and the respective year ranges was evaluated using Spearman’s rank correlation. The association between the number of plasmid replicons and phylogenetic groups was also evaluated using Spearman’s rank correlation. Statistical significance was defined as p□<□0.05.

To assess phage diversity in the isolate collection, VirSorter2 [72] (https://github.com/jiarong/VirSorter2) was used. For each *S. haemolyticus* genome analysed, the lengths of the identified phage DNA regions were summed to obtain the total phage DNA content per genome.

## Acknowledgments

We would like to acknowledge the Quadram Institute Sequencing team and Core Bioinformatics Team for all their help and expertise.

## Author Contributions

H.F, L.L. and M.W were involved in the study conception and design. Collection and processing of isolates was carried out by H.F., L.L., A.A-G., J.E.B., J.A.C., D.S., C.J.S., A.C.F., M.K. O’S., J.P.C., L.H. Both H.F. and L.L were involved with the data analysis and phenotyping of genomes and the first draft of the paper. H.F., L.L., J.P.C, L.H., A.C.F., W.v.S. and M.W were all involved in the production of the manuscript and scientific input.

## Ethical Statement

Isolates collected by the Quadram Institute from the Norfolk and Norwich University Hospital were from a prior study and were collected during routine surveillance. Clinical isolates were provided under NHS Research Ethics Committee approval to the Norwich Biorepository which banks blood, solid tissue, and bacterial isolates from the NNUH and research institutes on the Norwich Research Park, including the University of East Anglia, and makes these available to the research community.

The PEARL study has been reviewed and agreed by the Human Research Governance Committee at the Quadram Institute Bioscience and the London-Dulwich Research Ethics Committee (reference 18/LO/1703) and received written ethical approval by the Human Research Authority. IRAS project ID number 241880

## Supplementary Materials

**Supplementary Table 1 *S. haemolyticus* metadata for all genome sequences used in this study.**

Supplied as separate file

**Supplementary Table 2 A summary of AMR and virulence genes identified using Abricate (75% minimum DNA identify) from the CARD and VFDB databases [35, 68, 69].**

Supplied as separate file

**Supplementary Table 3** Scoary analysis of roary gene presence and absence in isolates from stool, blood, clinical sources. Also, neonatal blood isolates compared to adult blood isolates. Isolates with specificity > 50 and a best pairwise comparison p value of <0.0001 were classified as significant.

Supplied as separate file

**Supplementary Figure 1.**
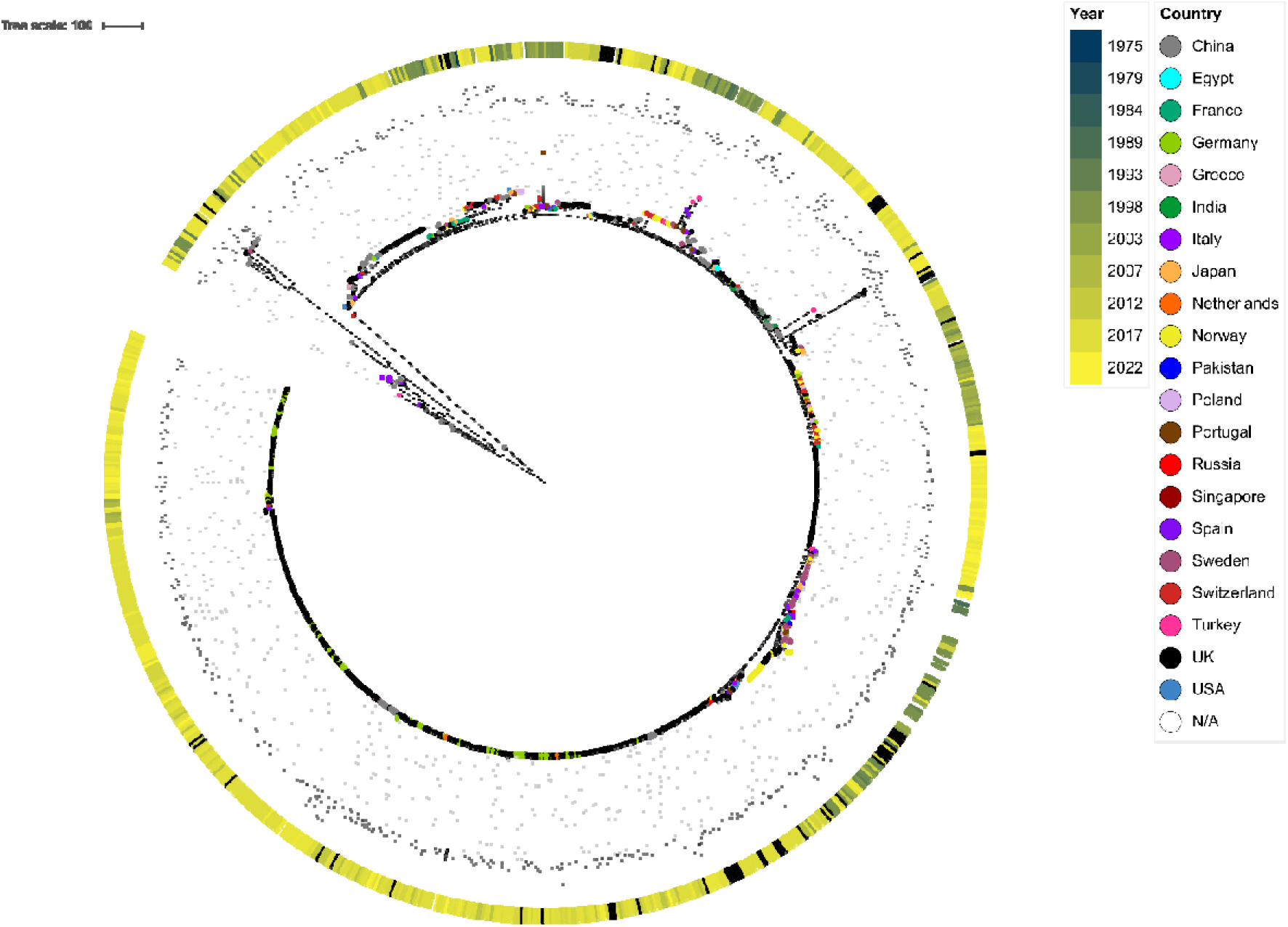
A phylogenetic tree based on the core SNPS of *S. haemolyticus*. Year of isolation is shown by coloured blocks in the outer circle and country of isolation by coloured labels for each isolate in the inner ring.

**Supplementary Figure 2.**
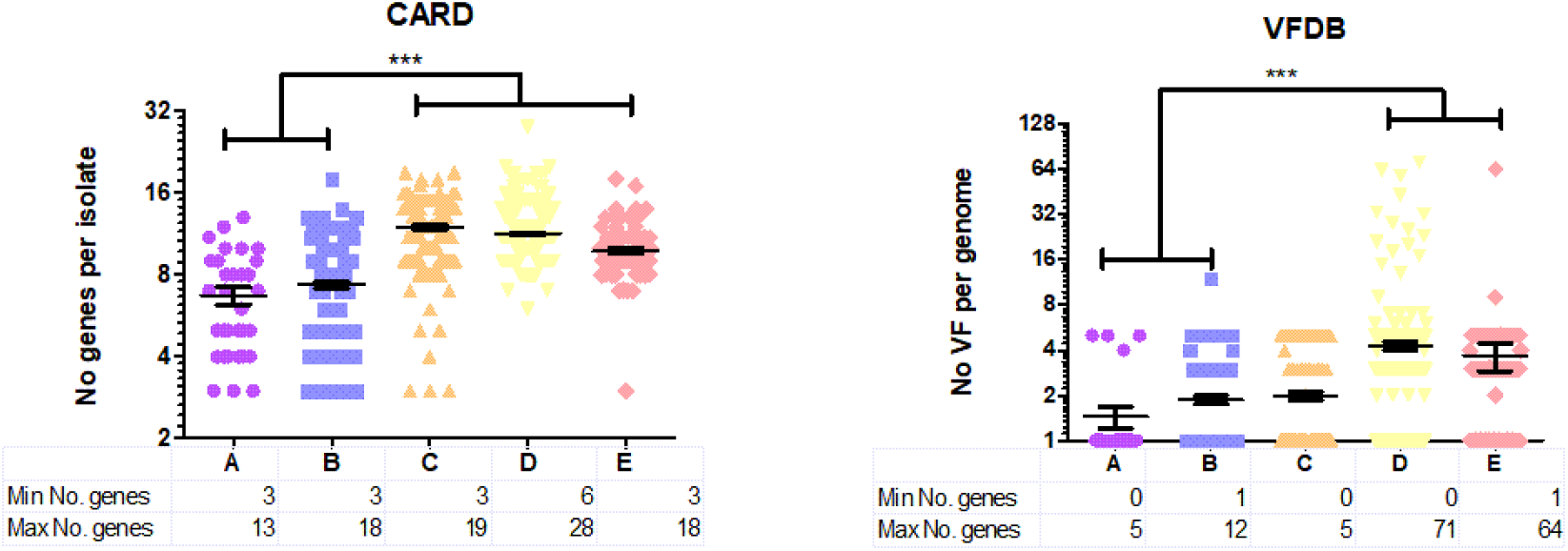
Numbers of AMR genes and virulence factors per isolate compared across the groups (Groups A-E). Numbers of genes was calculated by Abricate [68] using the CARD or VFDB databases [35, 69]. One way ANOVA was used to test statistical significance for differences between groups *** = p<0.05.

