## Supplementary figures and images for "*Staphylococcus haemolyticus* Population Genomics Provides Insights into Pathogenicity and Commensalism"

### Supplementary Figure 1

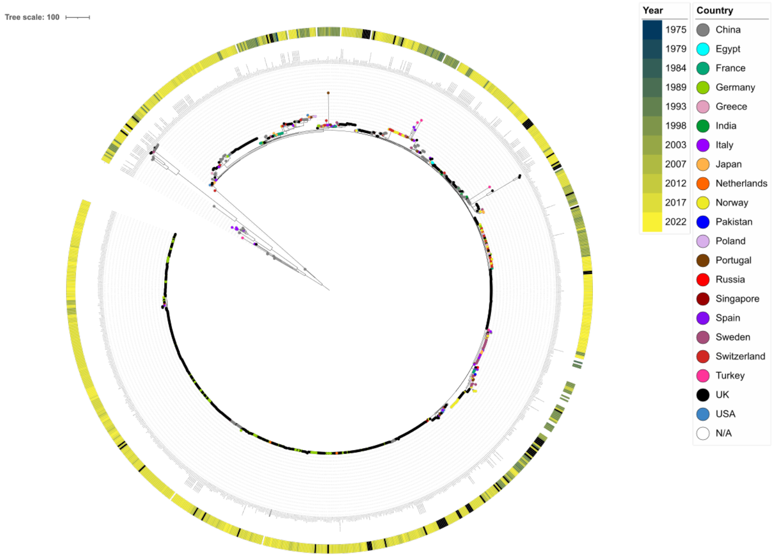

### Supplementary Figure 2

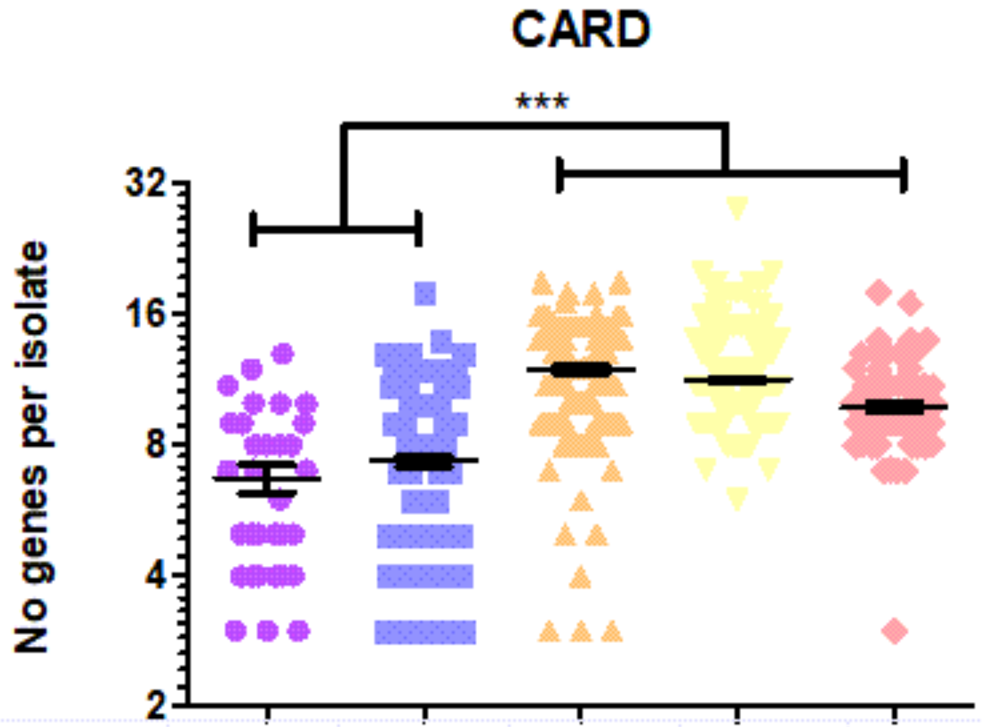


| | A | B | C | D | E |
|---|---|---|---|---|---|
| Min No. genes | 3 | 3 | 3 | 6 | 3 |
| Max No. genes | 13 | 18 | 19 | 28 | 18 |

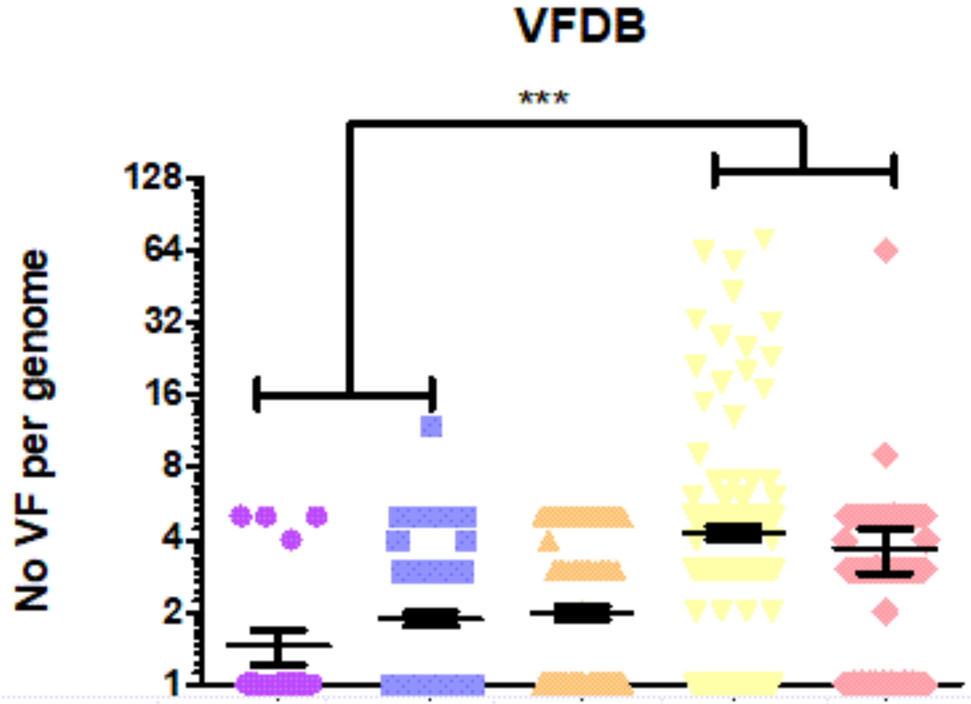


| | A | B | C | D | E |
|---|---|---|---|---|---|
| Min No. genes | 0 | 1 | 0 | 0 | 1 |
| Max No. genes | 5 | 12 | 5 | 71 | 64 |
